# Defective transcription in UPF1-deficient cells safeguards against replication stress induced transcription-replication collisions, driving drug resistance

**DOI:** 10.1101/2025.05.22.655556

**Authors:** Thomas Walne, Laura Maple, Lydia Ellis, Nan Li, Carl Smythe, Ruth Thompson

**Affiliations:** Division of Clinical Medicine, Faculty of Health, University of Sheffield, Sheffield. UK; School of Biosciences, Faculty of Science, University of Sheffield, Sheffield. UK; Nucleic Acids Institute, University of Sheffield, Sheffield. UK

**Keywords:** Mitosis, DNA damage, DNA replication, MiDAS, Runover replication, Transcription-replication conflicts

## Abstract

Accurate DNA replication is essential for the faithful transmission of genetic information to daughter cells. Disruption of this process leads to replication stress, which can trigger mutagenesis, double-stranded DNA breaks, and genomic instability. Transcription is a well-established source of replication stress, contributing through altered chromatin dynamics, DNA structural changes, and direct transcription–replication collisions (TRCs).

Here, we uncover a novel role for the RNA/DNA helicase UPF1 in maintaining replication fidelity and responding to replication stress. We show that cancer cells deficient in UPF1 exhibit elevated levels of spontaneous, transcription-dependent replication fork stalling and double-stranded breaks, along with heightened sensitivity to Rad51 and PARP1 inhibitors. Paradoxically, these cells also display resistance to exogenous replication stress, with reduced replication fork stalling, diminished mitotic delays, and decreased activation of mitotic DNA synthesis (MiDAS), a key salvage pathway under stress conditions.

Low UPF1 expression has been previously linked to drug resistance in renal carcinoma and correlates with poor prognosis across multiple cancer types. Our findings position UPF1 as a critical guardian against transcription-associated replication stress and suggest that UPF1 deficiency may underlie mechanisms of cancer therapy resistance whilst leaving cells vulnerable to the depletion of key DNA repair pathways. Targeting this vulnerability could offer a promising avenue for therapeutic intervention in UPF1-low tumours.

## Introduction

Up frameshift 1 (UPF1) was first identified and characterised as a helicase with a central role in mRNA decay pathways, the best characterised of which being nonsense mediated decay (NMD). Together with 6 other proteins (UPF2, UPF3, SMG1, SMG5, SMG6 and SMG7) UPF1 facilitates the selective destruction of newly synthesised mRNAs that possess a nonsense mutation [1]. In human cells, UPF1 also mediates other NMD independent RNA decay pathways including Staufen1-mediated decay and replication-dependent histone mediated decay [2, 3]. More recently, it has become clear that UPF1 plays a wider role in cellular RNA metabolism, regulating the stability and decay of a variety of specific RNAs including histone mRNA, long non-coding RNAs (lncRNAs) [4, 5] and microRNAs [6].

Aside from its canonical role in RNA surveillance, UPF1 has also been implicated in playing a major role in other cellular processes such as DNA replication [7], DNA repair [8] and cell cycle control [9, 10] however the precise mechanism of UPF1 in these pathways is unclear. Azzalin *et al* have shown that cells stably transfected with UPF1 shRNA experience a considerable replication defect, marked by an accumulation in early S phase and reduced BrdU incorporation. They also observed elevated PCNA on replication forks, indicative of replication fork stalling, as well as persistent activation of ATR and an increase in spontaneous levels of γH2AX in UPF1-deficient cells. Together these observations suggest that UPF1-deficient cells experience spontaneous replication fork collapse [7]. UPF1 has also been shown to interact with DNA polymerase delta [7, 11] further supporting a direct role in DNA replication. However, others have argued that the negative effects of UPF1 deficiency on replication could be due to defects in NMD altering gene expression [12]. Indeed, the role of UPF1 in the regulation of mRNAs has been linked to the DNA replication process previously, specifically through the controlled degradation of histone mRNAs. We previously identified a DNA-PK activated S phase checkpoint, which links histone transcription to DNA replication and leads to UPF1 activation [9, 10]. Interestingly, low levels of UPF1 have been detected in various cancers when compared to paired non-cancerous tissue and this down-regulation correlates with poor prognosis in these tumours [13, 14], however the reason for this is unknown.

In S phase of the cell cycle, transcription and replication occur simultaneously and as both use DNA as a template, careful coordination is required to avoid collisions between the machineries required for these two essential processes. These collisions are known as transcription-replication conflicts (TRCs) and are a major cause of genomic instability and DNA damage [15]. Recent single molecule studies which give direct visualisation of TRCs revealed post-replicative DNA:RNA hybrids behind replication forks following a TRC which resulted in replication fork slowing and reversal [16].

Replication stress during S-phase can result in a failure to complete DNA replication, leading to the emergence of under-replicated DNA regions following S-phase. Processing of this under-replicated DNA has been suggested to occur through distinct DNA replication processes in G2 [17, 18] and beyond into mitosis through mitotic DNA synthesis (MiDAS) [19–21] and G2 replication (termed G-mids). G-mids has been shown to occur at transcription start sites [18] and MiDAS is known to be caused by TRCs specifically in BRCA2-deficient cells [22].

UPF1 is a highly processive DNA:RNA helicase which has the ability to bind a multitude of DNA or RNA structures, including DNA:RNA hybrids possessing single strands of DNA or RNA with equal specificity, to catalyse the resolution of such structures [23].

Here we show an important role for UPF1 in DNA synthesis and the replication stress response. UPF1 deficiency through either siRNA mediated knockdown or CRISPR knockout leads to slowed replication speeds and elevated levels of replication fork stalling in unperturbed conditions. However, UPF1 deficient cells also exhibit high levels of resistance to replication stress inducing agents and simultaneously exhibit far lower levels of stalled replication forks than their wild type counterparts in response to these agents. Our data indicates that instead of acting on replication directly, UPF1 influences DNA replication stalling by regulating either transcription or the nascent RNA molecules and indeed we observe a transcriptional defect in these cells. Taken together these data present a possible mechanism for cancer drug resistance in UPF1 deficient cells along with a potential therapeutic angle for targeting these cells.

## Materials and Methods

### Cell Culture and Reagents

HeLa cells (ATCC) were cultured as previously described [24]. RPE UPF1^WT^ and RPE UPF^KO^ were a kind gift of Dr Greg Ngo. Where indicated, cells were treated with Carboplatin (Sigma-Aldrich), aphidicolin (Santa Cruz), RO3306 (Cayman Chemical), at indicated concentrations and with indicated length of time. Antibodies against UPF1 and RNAP II Ser2 were from Cell Signaling, γH2AX were from Novus Bio, and β-Tubulin were from Sigma. Appropriate secondary antibodies conjugated to horseradish peroxidase (DAKO) were used for the western blotting experiments, and Alexa-Fluor 488 and Alexa-Fluor 594 secondary antibodies (Invitrogen) were used for immunofluorescence microscopy.

### Flow cytometry

Cells were fixed in 70 % ethanol prior to staining. Following PBS washes to remove ethanol, cells were permeabilised by incubation in Flow Buffer 1 (0.5% BSA, 0.25% Triton-X). Flow Buffer 1 was removed after centrifugation and the cells were incubated in pH3 antibody (EMD Millipore, 3018868) diluted 1:100 in Flow Buffer 1 for 1.5 hours at room temperature. Cells were washed 3 times in Flow Buffer 2 (0.25% Triton-X) then resuspended in 100 µL Goat pAb anti-Rabbit IgG FITC secondary antibody, diluted 1:100 in Flow Buffer 1 and incubated for 30 minutes at room temperature in the dark. The cell pellets were resuspended in 400 µL PI (10 µg/µL stock solution in PBS) containing RNAse A (80μg/ml) and incubated for at least 30 minutes at 4 °C until processing.

The samples were processed on a FACSCalibur (BD Sciences) and analysed using Flowjo.

### RNAi and DNA transfection

siRNA transfections were performed using Dharmafect 1 siRNA transfection reagent (Horizon), DNA transfections using Lipofectamine 2000 (Thermo-Fisher Scientific) and siRNA/ DNA contranfections using Dharmafect Duo (Horizon) according to the manufacturer’s instructions. Cells were treated 48 hours post-transfection. siControl, siUPF1, siUPF2, siUPF3b, siSTAU1 siGenome SMARTpool siRNA pools were from Horizon Discovery.

### Western blotting

Cells were lysed using RIPA buffer and protein concentrations determined using a Bradford Assay. Lysates were separated by SDS-PAGE and transferred to nitocellulose. Blots were blocked in 5 % Milk in TBS (Tris-buffered saline) prior to overnight incubation with specified antibodies at 1:1000 in 5% milk diluted in TBS.

### Immunofluorescence microscopy

HeLa cells (5 x 10^4^) were seeded directly onto coverslips fixed in methanol or paraformaldehyde, permeabilised in 0.2% Triton-X100, blocked in 5% BSA and stained with the indicated antibodies. Alexa-Fluor 488 and Alexa-Fluor 594 secondary antibodies (Invitrogen) were used. In the final wash, cells were incubated with DAPI (Life technologies) and mounted to slides using Immumount (Thermo Fisher Scientific). Images were captured using a Nikon ECLIPSE Ti2 confocal microscope.

### Live cell Imaging

48 hours post transfection with indicated siRNAs, cells were trypsinised, exposed to ionising radiation (IR) whilst in suspension and reseeded to 24 well plates. Once adhered to the plates (3-4 hours post IR), the cells were loaded into imaging system, or following chemical administration as indicated. Live cell images were captured using ZEISS Cell discoverer 7 microscope every 5 minutes for a duration of 20 hours.

### Mitotic EdU incorporation assay

MiDAS assay was performed 48 hours post transfection as described by Garribba et al [25], and images were captured using Nikon ECLIPSE Ti2 confocal microscope.

### Clonogenic cell survival assay

UPF1 KO and UPF1 WT RPE cells were seeded at low density into 6-well plates and treated as indicated after a 3–5-hour incubation. The cells were then incubated for a further 7 days until sufficient colony growth was observed. Colonies were stained with methylene blue and colonies with ≥50 cells scored. Counts were then normalised for plating efficiency against the untreated control.

### Proximity Ligation Assay

RPE1 cells (3×10^6^) were seeded onto coverslips in a 6-well plate and left for 24h. Cells were then treated with or without aphidicolin for 23h, prior to fixation with ice-cold methanol at -20°C overnight. Cells were then blocked with 10% FBS in PBS at 37°C for 1h, before anti-POLR2A(pSer2) (NB100-1805, Novus, 1:400) and PCNA (PC10) sc-56 (Santa Cruz, 1:400) were added for 1h at 37°C for 1h. Proximity Ligation was then performed as described in the Duolink® In Situ Proximity Ligation Assay kit (Sigma). Coverslips were mounted using ProLong Gold with DAPI and imaged the following day using a Nikon Widefield microscope with 40x magnification.

## Results

### UPF1 is required for DNA damage-induced mitotic delay

We recently published a high-throughput screen for proteins required for a DNA damage-induced delay in mitotic progression [26]. In this screen, UPF1 knockdown resulted in a significant reduction in mitotic population following ionizing radiation. Here we show that siRNA-mediated depletion of UPF1 leads to the loss of DNA damage-induced mitotic delay (**Figure 1A**). UPF2, which binds to UPF1 and brings about a conformational change required for its activation [27, 28], is also required (**Figure 1A**). Whilst UPF2 interacts with UPF1 in a range of functions, siRNA-mediated depletion of UPF1 recruitment factors UPF3b and STAU1, which specifically recruit UPF1 to target mRNAs for NMD and SMD respectively, had no impact on mitotic progression (**Figure 1B**). This indicates that the function of UPF1 in mitotic progression is independent of NMD and SMD. Ectopic over-expression of a doxycycline inducible siRNA-resistant UPF1 was sufficient to rescue the effects of siRNA-mediated depletion of endogenous UPF1, demonstrating the change in phenotype is due to the specific loss of UPF1 (**Figure 1C**). Previous studies have demonstrated a role for ATR but not ATM in facilitating the chromatin recruitment of UPF1 for a seemingly NMD-independent function [7]. We found that whilst DNA damage induced mitotic transit delay does not require active ATM or ATR, small molecule inhibition of ATR is also sufficient to induce UPF1-dependent spontaneous transit delay (**Figure 1D**).

**Figure 1:**
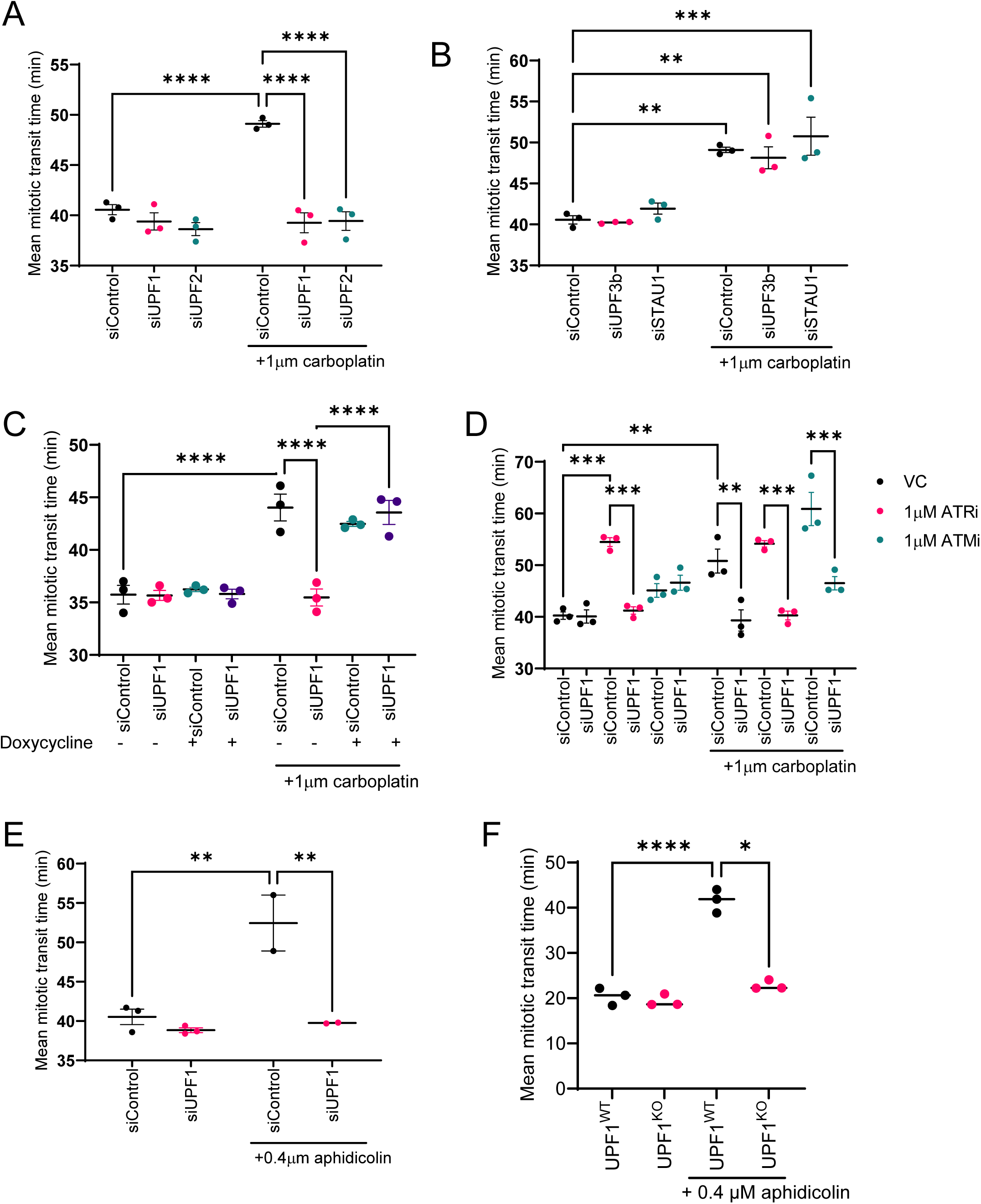
UPF1 and UPF2 are required for promoting a DNA damage-dependent mitotic delay. Time-lapse live cell microscopy analysis for the mean time taken to complete mitosis for 24 hours post treatment. **(A-B)** HeLa cells transfected with the indicated siRNAs and treated with 1µM Carboplatin as shown. **(C)** FLP-IN TReX UPF1^WT^ HeLa cells were treated as indicated with 1µg/ml Doxycycline, transfected with the indicated siRNAs for 48hrs. **(D)** Hela cells transfected with the indicated siRNAs and treated with 1µM ATRi or 1µM ATMi as indicated. **(E)** HeLa cells transfected with the indicated siRNAs treated with or without 0.4µM aphidicolin. **(F)** RPE UPF1^WT^ and UPF1^KO^ treated with or without 0.4µM aphidicolin. For all experiments 50 cells for each condition were counted and the data illustrated represents the overall mean of each independent experiment +/- SEM (N=3). One way ANOVA with Dunnett’s correction test for multiple comparisons was performed to determine statistical significance. (* denotes p ≤ 0.05, ** denotes p ≤ 0.01, *** denotes p ≤ 0.001 and **** denotes p ≤ 0.0001). *Ctrl = Control, VC = Vehicle Control, Dox = Doxycycline*

We have previously found that replication stress induced by low dose aphidicolin, also induces mitotic transit delay [26]. This delay was found also to be UPF1-dependent (**Figure 1E**). As further confirmation that UPF1 is required for mitotic transit delay, we scored mitotic duration in paired RPE1 UPF1^WT^ and UPF1^KO^ cells in response to aphidicolin and found that the knockout cells did not exhibit mitotic delay in response to UPF1 (**Figure 1F).**

### UPF1 depletion results in reduced MiDAS and G2 synthesis under conditions of replication stress

Whilst it is unclear why cells undergo a delayed mitotic transit in response to DNA damage and replication stress, work from our lab and others has revealed a close link between mitotic duration and the occurrence of MiDAS in prophase [29–31]. UPF1 is known to interact with POLD3 and has been implicated in R-loop metabolism, both of which are associated with MiDAS. Thus, we chose to investigate the effects of UPF1 depletion on MiDAS. Using standard conditions to induce MiDAS (**Figure 2A**), we found that cells depleted for UPF1 using either siRNA (**Figure 2B-C**) or CRISPR (**Figure 2D-E**), exhibited far fewer EdU foci in prophase than their control counterparts, indicating a reduction in mitotic DNA synthesis under these conditions. FANCD2 is common marker of replication stress and known to colocalise with or mark common fragile sites that will undergo MiDAS in prophase [32]. We observed a significant reduction in FANCD2 foci in UPF1-deficient cells treated with low-dose aphidicolin (**Figure 2E-F**). Taken together, these data suggests that the reduction in MiDAS in these cells following UPF1 depletion is due to the relative absence of under-replicated regions of DNA prior to mitotic entry, rather than a failure to undertake MiDAS, since a defective MiDAS pathway would not reduce the frequency of FANCD2 foci in prophase cells [33].

**Figure 2:**
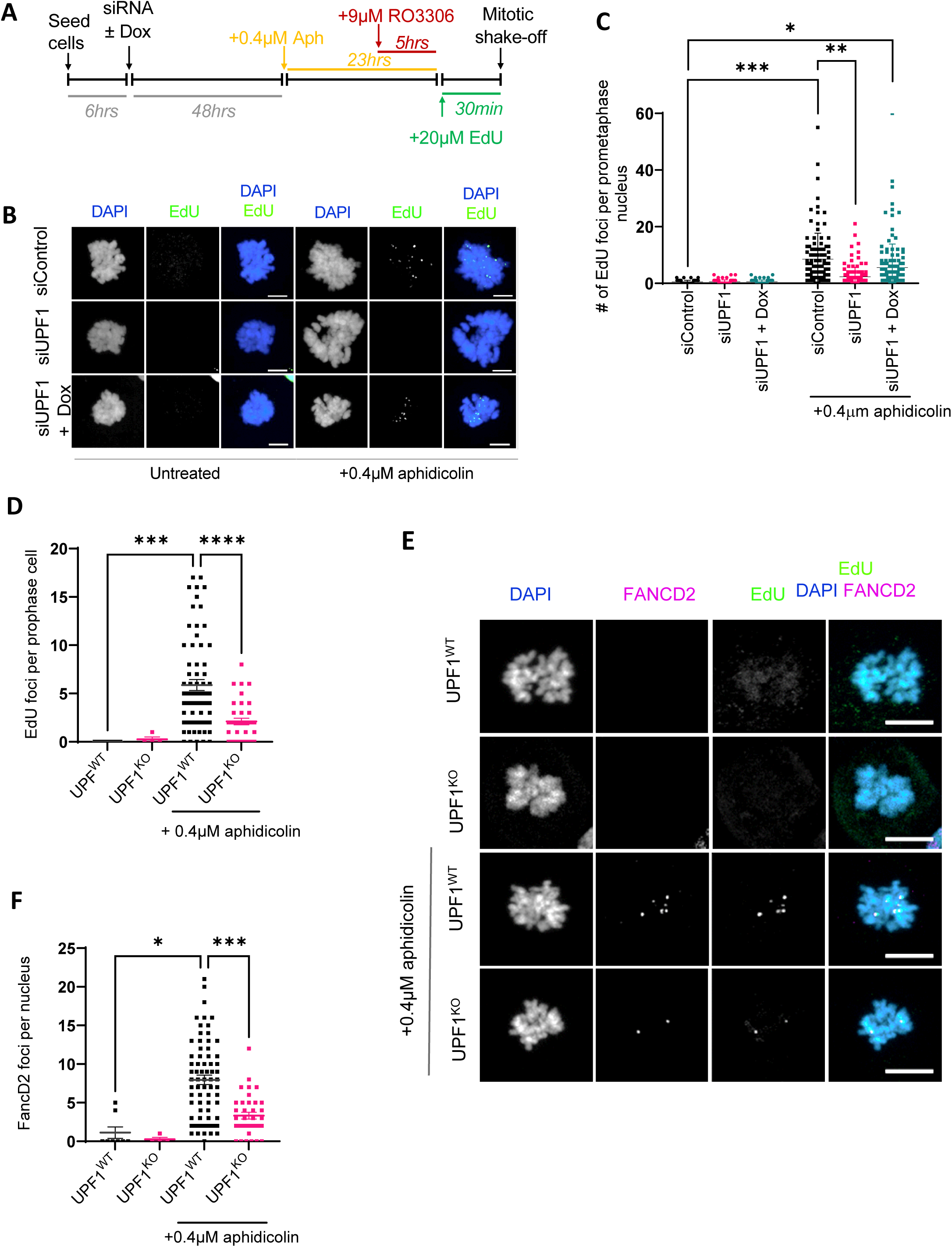
UPF1 is required for Mitotic DNA synthesis following mild replication stress. **(A)** Schematic depicting experimental outline. Cells were transfected with indicated siRNAs and induced with 1µg/ml doxycycline as described. Cells were treated with 0.4µM Aphidicolin as indicated for 23hrs prior to mitotic shake-off and incubation with 20μM EdU for 25mins prior to fixation. **(B)** Representative immunofluorescence images showing EdU and DAPI of prophase cells. **(C)** Quantification of (B) FLP-IN TReX UPF1^WT^ HeLa cells were treated as indicated with 1µg/ml Doxycycline, transfected with the indicated siRNAs for 48hrs. EdU foci per prometaphase cell. 50 cells for each condition were quantified and the data illustrated represents individual EdU foci from three biological replicates. N=150 **(D)** RPE UPF1^WT^ and UPF1^KO^ cells were treated with 0.4µM Aphidicolin as indicated for 23hrs prior to mitotic shake-off and incubation with 20μM EdU for 25mins prior to fixation. Quantification of EdU foci per prometaphase cell. **(E)** Representative images of D and **(F)** Quantification of FancD2 foci in UPF1^WT^ and UPF1^KO^ treated with or without 0.4µM aphidicolin. 50 cells for each condition were quantified and the data illustrated represents individual EdU foci from three biological replicates n=150. The error bars represent the mean +/- SD. One way ANOVA with Dunnett’s correction test for multiple comparisons was performed to determine statistical significance (* denotes p ≤ 0.05, ** denotes p ≤ 0.01 and *** denotes p ≤ 0.001). *Dox = Doxycycline, Aph = Aphidicolin*

MiDAS was first identified in cells exposed to mild replication stress and was proposed as a mechanism by which cells entering mitosis can restart DNA synthesis to complete replication of under-replicated or difficult-to-replicate loci known as chromosome fragile sites [20, 21, 33]. In all these studies, the CDK1 inhibitor RO3306 was used to arrest cells at the G2/M boundary in order to enhance the mitotic population for analysis. More recently it has been demonstrated that the phenomenon of replication restart in mitosis is brought about through off-target effects of RO3306 on CDK2 [17]. It is suggested that DNA synthesis does not in fact stop at the end of S phase, and under conditions of mild replication stress, continues into G2 and beyond into mitosis [17].

Furthermore, DNA synthesis detected in G2, in the absence of RO3306, appears to be independent of several canonical MiDAS proteins including MUS81 and POLD3. To determine whether the involvement of UPF1 in MiDAS was due to forced replication restart brought about by RO3306 treatment, we assessed DNA replication dynamics throughout the cell cycle in asynchronous conditions in our paired UPF1^WT^ and UPF1 ^KO^ cells (**Figure 3**). Following treatment with aphidicolin, cells were pulse-labelled for 30 minutes with EdU, then harvested and stained for EdU and FANCD2 (**Figure 3A)**. As with the mitotic DNA synthesis observed in **Figure 2**, there was a significant reduction in replication stress-induced EdU and FANCD2 foci in non-S phase cells in the absence of UPF1 (**Figure 3B-D**). Interestingly however, there was also a reduction in overall EdU intensity in S phase cells in the absence of UPF1 (**Figure 3**E-F),suggesting that UPF1 loss may affect global DNA replication rates or efficiency of origin firing. Our data also shows that the UPF1 ^KO^ cells have slightly elevated levels of FANCD2 foci in S and G2 in untreated conditions indicating that UPF1 loss on its own may lead to replication stress (**Figure 3E, G**). Together this indicates that whilst loss of UPF1 promotes replication stress in unperturbed cells, it also protects against aphidicolin-induced replication stress.

**Figure 3:**
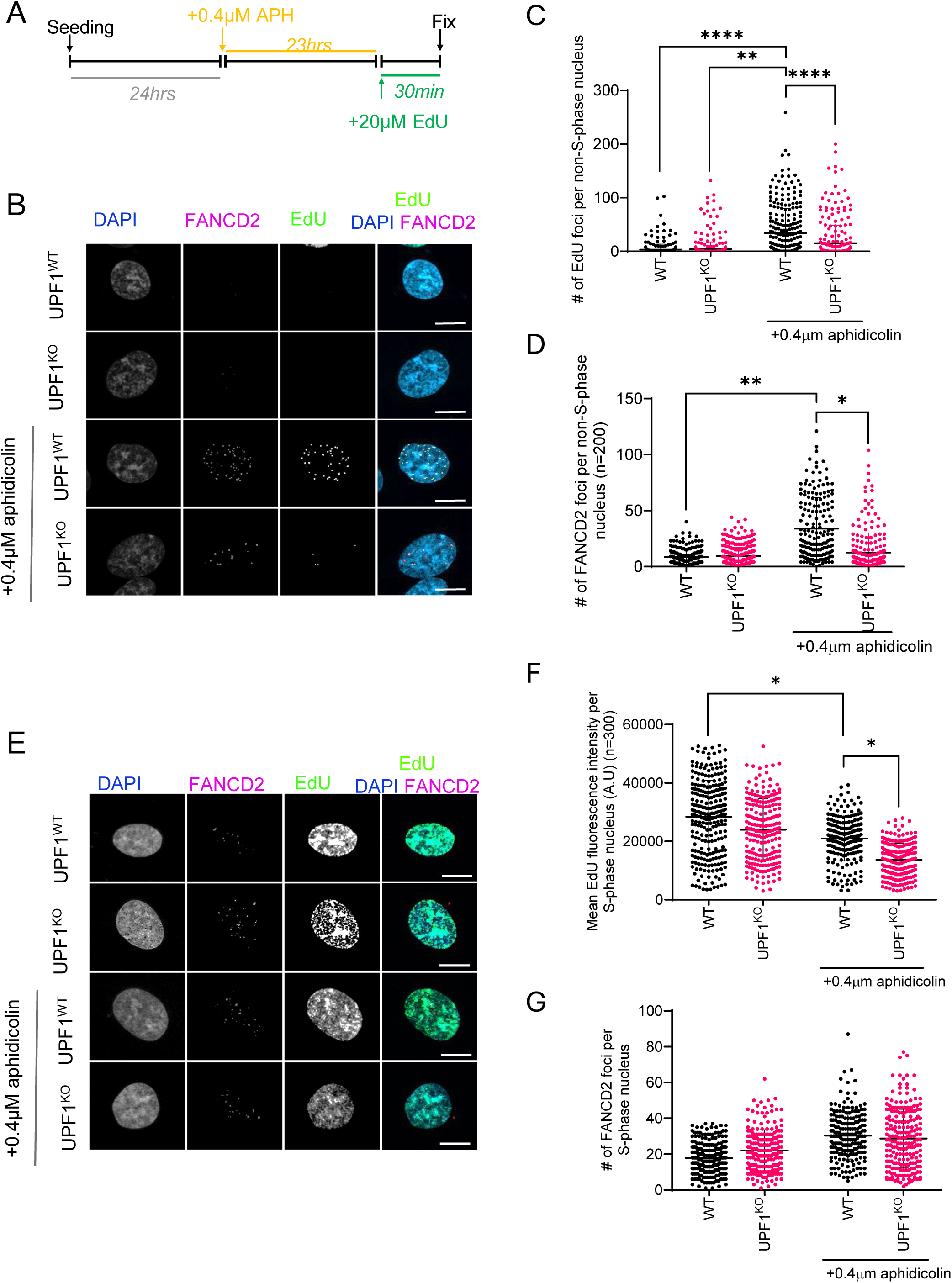
UPF1 is required for continuous DNA synthesis outside of S phase following mild replication stress. **(A)** Schematic depicting experimental outline. RPE UPF^WT^ and UPF1^KO^ cells were seeded to coverslips, treated with 0.4µM Aphidicolin as indicated for 23hrs prior to incubation with 20μM EdU for 25mins prior to fixation. Coverslips were stained for EdU, and FancD2. Representative immunofluorescence images for RPE^WT^ cells at each stage of the cell cycle untreated and with 0.4µM aphidicolin for 24hrs, stained for EdU, FANCD2, H3PS10 and DAPI. **(B)** Representative images for non-S-phase cells. **(C-D)** Quantification of nuclear EdU and FANCD2 foci per non-S-phase cell respectively. **(E)** Representative images for S-phase cells. **(F-G)** Quantification of mean nuclear EdU intensity and FANCD2 foci per S-phase cell respectively. Data illustrated are a representative graph taken from one biological replicate. For EdU foci, intensity and FANCD2 foci in all cell cycle phases a One-way ANOVA with Dunnett’s correction test for multiple comparisons was performed on the mean nuclear foci from each biological replicate to determine statistical significance (N=3) (* denotes p ≤ 0.05, ** denotes p ≤ 0.01, *** denotes p ≤ 0.001, **** denotes p ≤ 0.0001).

### UPF1 deficiency results in slowed DNA replication and collapsed replication forks in unperturbed conditions but rescues stalled replication forks induced by aphidicolin

Azzalin et al have suggested that UPF1 is required for S phase progression, showing that cells lacking UPF1 appear to accumulate in early S phase [7]. UPF1^KO^ cells proliferate more slowly than their wild type counterparts [8] which also suggests a possible replication defect. We therefore performed DNA fibre analysis to fully assess the effect of UPF1-deficiency on DNA replication. DNA fibre analysis revealed significant reductions in replication fork speed (**Figure 4A**) and increased levels of replication fork stalling as determined by reduced sister fork symmetry (**Figure 4B**) in siUPF1-treated cells compared to the siControl-treated cells in unperturbed conditions. Together this suggests that loss of UPF1 leads to the spontaneous stalling of replication forks. In control cells, as expected, treatment with low-dose aphidicolin resulted in replication fork stalling and significantly reduced fork speeds. Consistent with our FANCD2 data (**Figure 3G**), however, whilst aphidicolin treatment still reduced replication speeds in UPF1-deficient cells (**Figure 4A**), there was a significant increase in sister fork symmetry suggesting reduced replication fork stalling under these conditions (**Figure 4B**). This effect was further replicated in the paired RPE UPF1^WT^ and UPF1^KO^ cell lines (**Figure 4C-D**). Since UPF1 is an RNA/DNA helicase whose function impacts directly on transcription [34], it is possible that defective transcriptional processes in the absence of UPF1 are responsible for promoting replication fork stalling. To test this, we hypothesised that inhibiting transcription with short-term exposure to the transcription inhibitor 5,6-dichloro-1-β-D-ribofuranosylbenzimidazole (DRB) would rescue spontaneous replication fork stalling. Indeed, the slowed replication speeds and loss of sister fork symmetry in UPF1-deficient cells were restored to WT levels following treatment with DRB (**Figure 4C-D**). In a similar manner, DRB treatment was also able to rescue spontaneous DNA damage (**Figure 4E-F**) which has been reported previously in cells depleted for UPF1 [7]. Together this indicates that the effect of UPF1-deficiency on replication fork stalling is dependent on active transcription.

**Figure 4:**
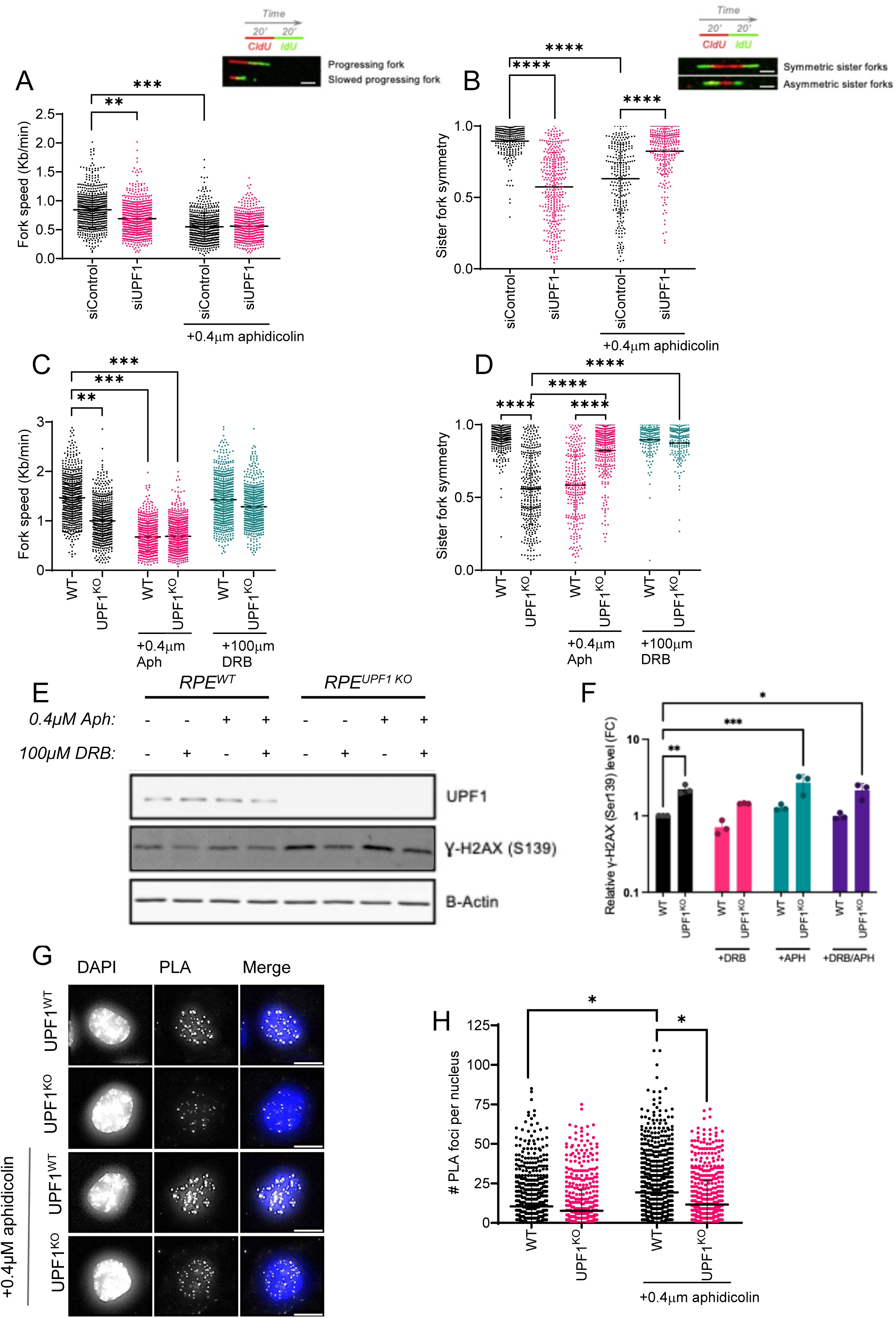
Loss of UPF1 promotes spontaneous replication fork stalling. **(A)** HeLa cells were treated with the indicated siRNAs for 48 hours then treated without and with 0.4µM aphidicolin for 24hrs. Cells were sequentially pulse treated for 20 minutes with CldU and IdU then the DNA fibres were combed onto coverslips and analysed by fluorescence microscopy. Quantification of progressing fork speeds. **(B)** HeLa cells were treated with the indicated siRNAs for 48 hours then treated without and with 0.4µM aphidicolin for 24hrs. Quantification of sister fork symmetry. **(C)** UPF1^WT^ and UPF^1^ ^KO^ RPE1 cells treated without and with 0.4µM aphidicolin for 24hrs in the presence and absence of DRB. Quantification of progressing fork speeds. **(D)** UPF1^WT^ and UPF^1^ ^KO^ RPE1 cells treated without and with 0.4µM aphidicolin for 24hrs in the presence and absence of DRB. Quantification of progressing fork speeds. A minimum of 200 individual DNA fibres were analysed per condition and a One-way ANOVA with Dunnett’s correction test for multiple comparisons was performed on the mean of each biological replicate to determine statistical significance (* denotes p ≤ 0.05, ** denotes p ≤ 0.01, *** denotes p ≤ 0.001, **** denotes p ≤ 0.0001). **(E)** UPF1^WT^ and UPF^1^ ^KO^ RPE1 cells treated without and with 0.4µM aphidicolin for 24hrs were lysed and analysed by western blot. Blot is representative data of 3 independent experiments. **(F)** densitometric analysis of E. γH2AX values shown normalised against GAPDH. **(G)** Representative images for **(H)** UPF1^WT^ and UPF^1^ ^KO^ RPE1 cells treated without and with 0.4µM aphidicolin for 24 hrs had the PLA assay performed. 50 cells per condition were score for number of foci and mean fluorescent intensity (N=3). One-way ANOVA test with Dunnett’s correction test for multiple comparisons was carried out on the mean of each biological replicate to determine statistical significance (ns denotes p > 0.5 and ** denotes p ≤ 0.01).

Having shown that UPF1-deficiency promotes transcription-dependent replication fork stalling, slowed fork progression and DNA damage, we wanted to test whether loss of UPF1 affected the occurrence of transcription-replication collisions (TRCs) both in unperturbed and mild replication stress conditions. To assess this, we performed proximity ligation assays (PLA) between PCNA and Ser2 phosphorylated-RNAPII-CTD (**Figure 4G-H**). As expected, aphidicolin induced a significant increase in the number of PLA foci in UPF1^WT^ cells which was not observed in the UPF1^KO^ cells, suggesting that loss of UPF1 does indeed reduce the occurrence of TRCs following mild replicative stress. However, we did not observe any increase in PLA foci in the untreated UPF1^KO^ cells when compared with the UPF1^WT^. It is possible that transcription-dependent replication fork stalling that occurs in the absence of UPF1 occurs independently of RNAPII or more specifically independent of Ser2 phosphorylated RNAPII.

### UPF1-deficient cells exhibit a transcriptional defect

UPF1 has previously been shown to associate with gene coding DNA regions and correlate with RNAPII-mediated transcription and RNA levels [35], and to play a role in the release of newly transcribed RNA molecules from the gene locus [36]. Since we have demonstrated that loss of UPF1 results in transcription dependent genomic stability we wished to determine whether there is indeed any disruption to transcription in the absence of UPF1. We assessed the level of Ser2 phosphorylated RNAPII-CTD, a marker for active transcription elongation of which UPF1-deficient cells display significant reductions in by both western blot (**Figure 5A-B**) and immunofluorescence microscopy (**Figure 5C**). Interestingly, whilst we observed significant reductions in Ser2 phosphorylated RNAPII we detected no changes in Ser5 phosphorylated RNAPII levels (**Figure 5D)**, a marker for initiating transcription or total RNAPII (**Figure 5E)**. We also assessed nascent RNA production through incubation with 5-ethynyl uridine (5-EU), a modified uridine nucleoside which is actively incorporated into nascent mRNA and can be detected using Click-iT labelling. UPF1-deficient cells display a 25% reduction in nascent mRNA production (**Figure 5F-G**), suggesting that in these cells there is a significant defect in productive transcription, whilst no observable changes in transcriptional initiation. Together these data support the existence of a transcriptional defect in UPF1-deficient cells which predisposes them to DNA replication stress and DNA damage.

**Figure 5:**
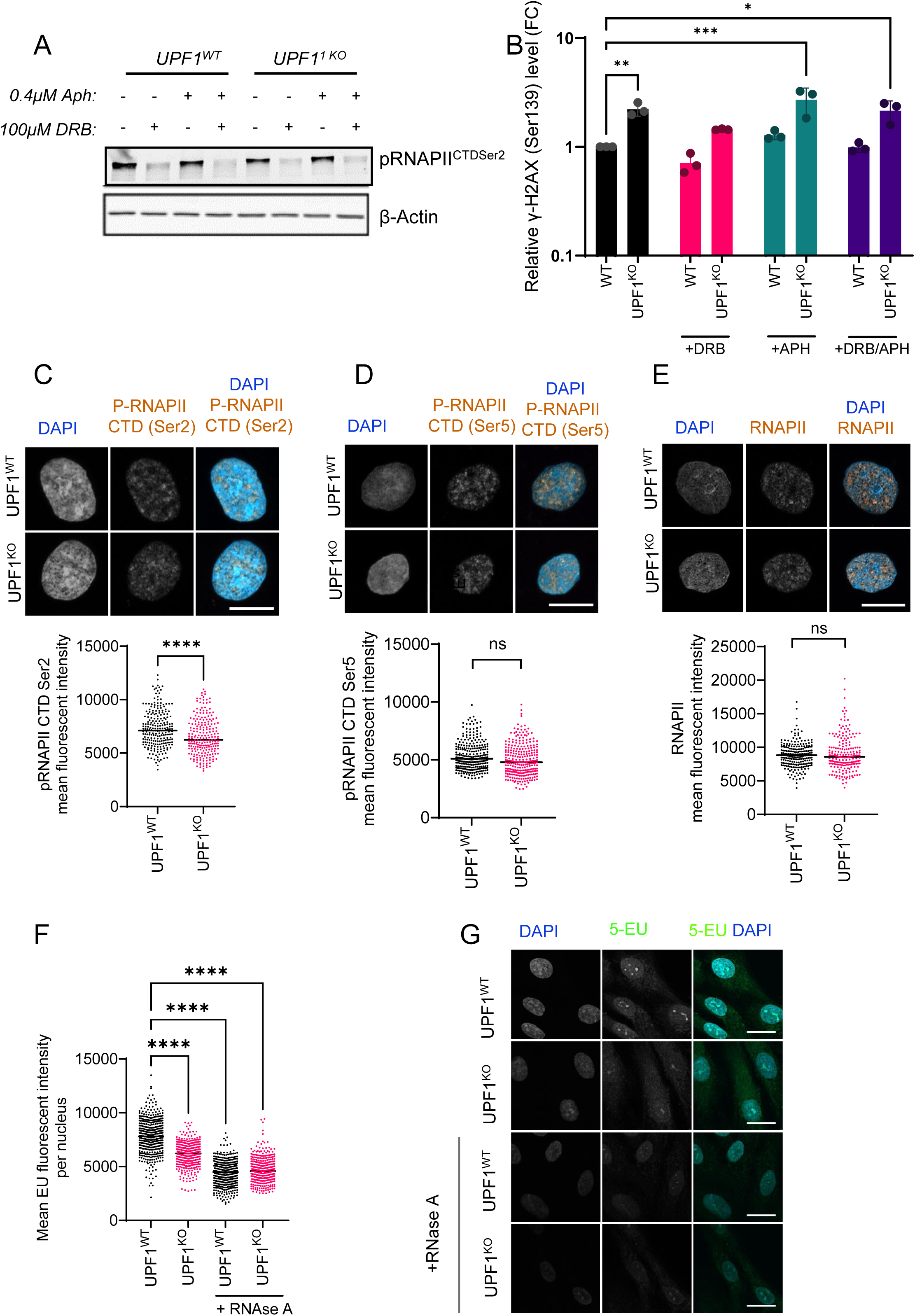
Loss of UPF1 leads to reduced levels of replication stress-induced R-loops and TRCs. **(A)** UPF1^WT^ and UPF^1^ ^KO^ RPE1 cells treated without and with 0.4µM aphidicolin for 24 hrs were stained for s9.6 and analysed by fluorescence microscopy. 50 cells were scored per condition. One-way ANOVA test with Dunnett’s correction test for multiple comparisons was carried out on the mean of each biological replicate to determine statistical significance. **(B)** Representative images of (A). **(C)** Representative images for **(D)** UPF1^WT^ and UPF^1^ ^KO^ RPE1 cells treated without and with 0.4µM aphidicolin for 24 hrs had the PLA assay performed. 50 cells per condition were score for number of foci and mean fluorescent intensity (N=3). One-way ANOVA test with Dunnett’s correction test for multiple comparisons was carried out on the mean of each biological replicate to determine statistical significance (ns denotes p > 0.5 and ** denotes p ≤ 0.01).

### UPF1-deficient cells are resistant to replication stress inducing agents but exhibit hypersensitivity to inhibitors of proteins required for homologous recombination and single strand break repair

We have found that whilst UPF1-deficient cells experience high levels of endogenous replication stress as a result of defective transcriptional processes, they do not show evidence of TRCs or experience replication fork stalling in response to mild replicative stress induced by low dose aphidicolin. These data suggest that UPF1-deficient cells would be resistant to chemotherapeutic agents which induce replication stress. It has previously been observed that low levels of UPF1 is a poor prognostic marker for renal clear cell carcinoma [13] and dysregulation of the UPF1 signalling axis through loss of UPF2 or SMG8:SMG9 mediates ATRi resistance in gastric cancer lines [37, 38]. We have observed that cells deficient for UPF1 are indeed more resistance to aphidicolin compared to their WT counterparts (**Figure 6A**), an effect which is more significant when cells are allowed to recover from a pulse of aphidicolin for 72hrs (**Figure 6B**). We also observed increase resistance to carboplatin (**Figure 6C**). The elevated endogenous levels of DNA breaks and stalled replication forks in UPF1^KO^ cells led us to hypothesise that UPF1-deficient cells, whilst exhibiting resistance to replication stress-inducing agents, may be sensitive to inhibitors of DNA repair pathways. We found this to be the case, with the UPF1^KO^ cells exhibiting increased sensitivity to Rad51 inhibitors at concentrations which had little effect on the wild type cells (**Figure 6D**). These cells were also found to be more sensitive to PARP inhibition (**Figure 6E**).

**Figure 6.**
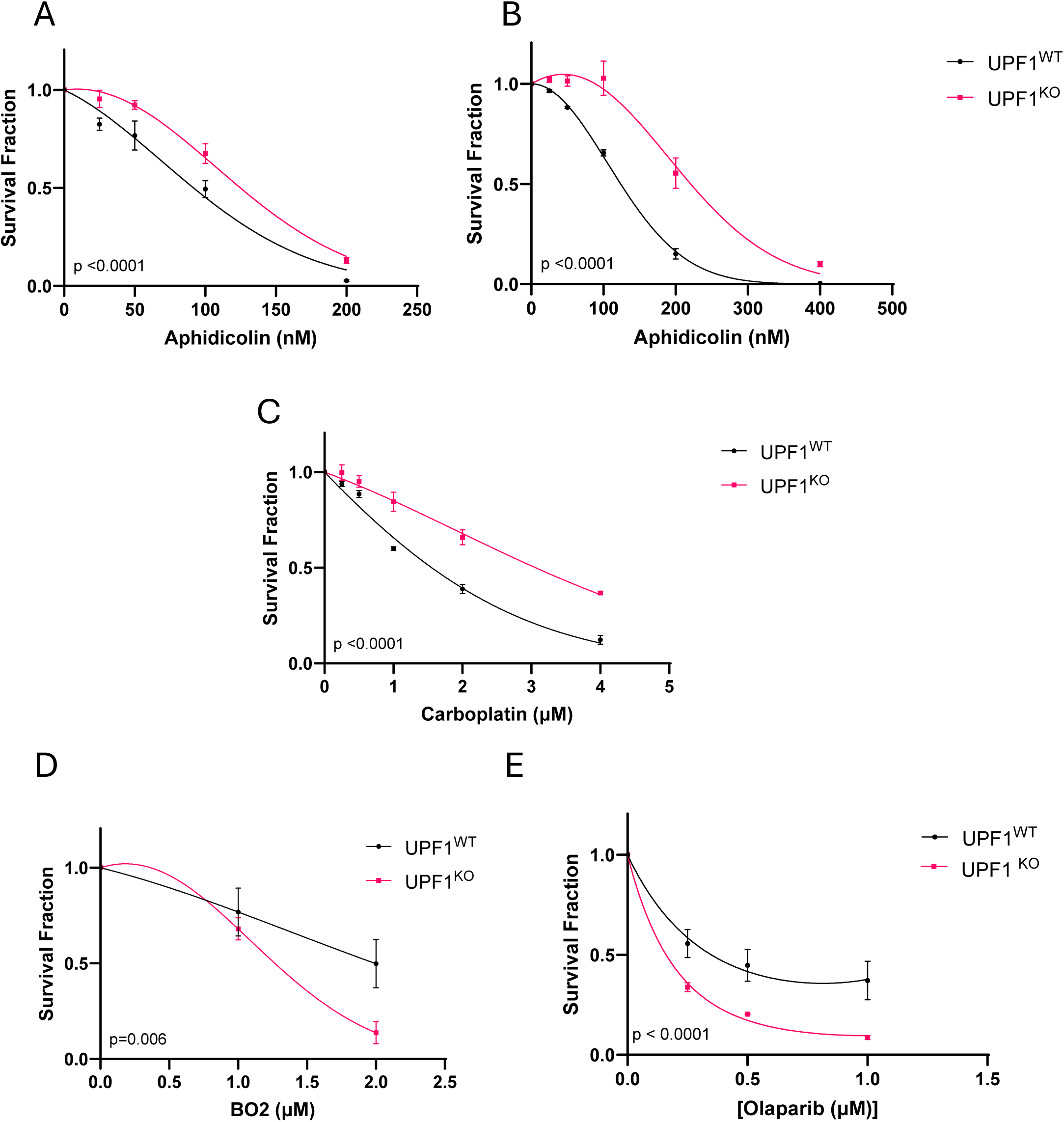
UPF1 deficiency confers resistance to replication stress but sensitivity to Rad51 and PARP1 inhibitors. **(A)** UPF1^WT^ and UPF^1^ ^KO^ RPE1 cells were plated at low density, treated as indicated and allowed to form colonies for 7 days. Colonies were then counted and survival calculated according to plating efficiency of cell line. Each assay was performed in triplicate. Error bars represent mean +/- SEM of three biological replicates. Non-linear regression comparison was carried out on the survival fraction using the Extra sum-of-squares F test. **(B)** UPF1^WT^ and UPF^1^ ^KO^ RPE1 cells were plated at low density, treated as indicated for 72 hours, washed three times and allowed to form colonies for 7 days. Colonies were then counted and survival calculated according to plating efficiency of cell line. Each assay was performed in triplicate. Error bars represent mean +/- SEM of three biological replicates. Non-linear regression comparison was carried out on the survival fraction using the Extra sum-of-squares F test. **(C-E)** UPF1^WT^ and UPF^1^ ^KO^ RPE1 cells were plated at low density, treated as indicated and allowed to form colonies for 7 days. Colonies were then counted and survival calculated according to plating efficiency of cell line. Each assay was performed in triplicate. Error bars represent mean +/- SEM of three biological replicates. Non-linear regression comparison was carried out on the survival fraction using the Extra sum-of-squares F test.

Taken together we have characterised a novel role for UPF1 in safeguarding DNA replication progression in unperturbed cell cycles through the maintenance of effective transcription. UPF1-deficient cells also demonstrate significant alterations in their response to exogenous replication stress resulting in cancer drug resistance and providing an explanation for poor prognosis in patients with UPF1-deficient tumours.

## Discussion

UPF1 has long been studied for its role in RNA surveillance, coordinating well characterised mRNA decay pathways such as NMD, SMD and replication-dependent HMD; and the effects that dysregulation of these pathways have on cancer cell biology, including increased stemness and chemotherapeutic resistance [4, 39–41]. Beyond these functions, UPF1 has been suggested to have more direct roles in both DNA replication and repair [7, 8], which would make it an interesting target for anti-cancer therapies, however little is known about this role. Reduced UPF1 expression has been documented in several cancers including hepatocellular cancer [5, 42, 43], gastric cancer [44], lung adenocarcinoma [45, 46], pancreatic cancer [47] and ccRCC where is has been shown to correlate with poor patient prognosis and increased invasion, migration and proliferation. However, the reasons for this are unclear.

Here we show how loss of UPF1, through siRNA-mediated depletion or CRISPR-Cas9 mediated knockout, alters the DNA replication stress response in both cancerous and untransformed human cell lines, resulting in the loss of replication stress-induced phenotypes including mitotic delay, MiDAS and replication fork stalling in S-phase. The reduction of these phenotypes in UPF1-deficient cells ultimately drives resistance to replication-targeting chemotherapeutics, providing a potential explanation for why low UPF1 expression correlates with poor patient prognosis [13].

UPF1 has been shown to interact with the regulatory subunit of DNA polymerase delta p66 and as a result has previously been implicated in both global and specific DNA replication processes in telomeric regions [7] and a slower cell cycle than their wild-type counterparts [8]. Here, we demonstrate that loss or depletion of UPF1 results in reduced replication fork speeds and increased levels of spontaneous replication fork stalling in an otherwise unperturbed cell cycle. Interestingly however, UPF1-deficiency was found to rescue replication fork stalling induced by the DNA polymerase inhibitor aphidicolin. This, taken together with previous literature demonstrating a key role for UPF1 in transcription through RNAPII progression [35, 48], release of nascent mRNA [36] and subsequent dissociation of the transcriptional machinery [35], led us to question whether the adverse effects observed in the absence of UPF1 on DNA replication may actually be via transcription. Indeed, we have observed significant disruption to transcription in the absence of UPF1, including a marked reduction in Ser2 phosphorylated RNAPII-CTD, a marker of active transcriptional elongation as well as a 25% reduction in nascent mRNA production. We show that acute inhibition of RNAPII-mediated transcription, through treatment of a high dose of the CDK9 inhibitor DRB, is sufficient to rescue spontaneous replication fork stalling and DNA damage caused by UPF1 deficiency. This suggests that defective RNAPII elongation and subsequent RNAPII stalling in the absence of UPF1 underlies spontaneous replication stress and DNA damage, since DRB treatment results in promoter-proximal paused RNAPII, prior to the initiation of transcription elongation [49].

Consistent with this hypothesis, as previously observed in UPF2 or SMG8:SMG9 deficient cells [37, 38] a proximity ligation assay between RNAPII-CTD phosphorylated at Serine 2 and PCNA, to assess the occurrence of TRCs, showed that UPF1-deficient cells exhibited significantly less PLA foci following treatment with aphidicolin when compared to the wild-type counterparts. Slowed or defective transcription elongation in the absence of UPF1 would explain why there is an observed reduction in frequency of TRCs following mild replicative stress, which results in the slowing of the DNA replicative machinery, therefore reducing the occurrence of such collisions.

Since we do not observe a spontaneous increase in TRCs in the absence of UPF1 by our PLA assay, it is possible that replication fork stalling during an unperturbed cell cycle occurs independently of RNAPII. We have observed a global reduction in RNAPII Ser2 phosphorylation in both untreated and aphidicolin treated conditions by western blotting and immunofluorescence potentially explaining the apparent reduction in TRCs. Instead, we propose that collisions between the replisome and bound nascent mRNAs are responsible for fork collision events.

Taken together we propose that UPF1 drives effective transcription and release of nascent RNA molecules, which in turn contribute to the maintenance of the transcription: replication balance. In control cells, aphidicolin reduces replication speed resulting in TRCs through the transcription machinery colliding with the stalled or slowed replication machinery, however in the absence of UPF1, the slowed transcription machinery compensates for the slowed replication machinery, meaning there are fewer TRCs and replication is able to continue (**Figure 7).**

**Figure 7:**
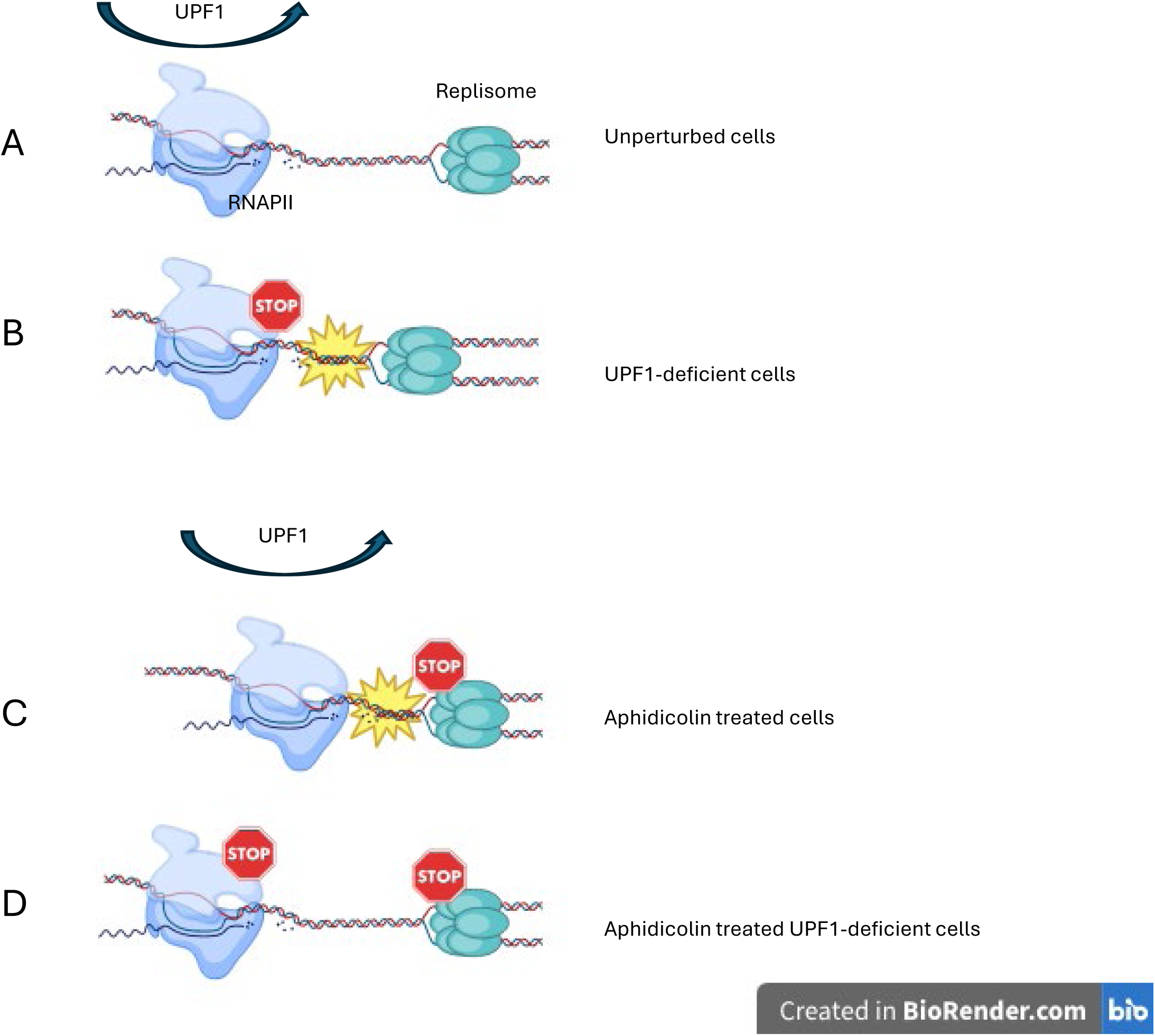
UPF1 model. **(A)** In normal cells, transcription and replication exist in S phase in a careful balance meaning there are no collisions or conflict. **(B)** UPF1 deficiency leads to an accumulation of R-loops which results in slowing of the transcription machinery causing the replication machinery to collide with it. **(C)** In normal cells, replication-stress inducing agents such as aphidicolin, slow the replication machinery, causing the transcription machinery to collide with it. **(D)** When replication-stress inducing agents are used in UPF1-deficient cells, both transcription and replication are slowed meaning they are once again in balance.

We show here for the first time that UPF1-deficiency results in a transcriptional defect which leads to slowed DNA replication and increased levels of replication fork stalling. In turn, this defect protects the cells from exogenous replication stress resulting in cancer drug resistance but results in hypersensitivity to inhibitors of DNA repair. Taken together these data provide an explanation for poorer prognosis in cancer patients with UPF1 deficient tumours whilst also discovering an alternative therapeutic vulnerability of these tumours.

## Acknowledgements

The authors would like to thank Darren Robinson for microscopy advice, Helen Bryant and Spencer Collis for helpful discussion and technical advice and Helen Bryant and Greg Ngo for the sharing of reagents. Thomas Walne acknowledges The Humane Research Trust for funding his PhD through The Ken Cholerton Studentship Award (RT) and Laura Maple acknowledges EPSRC for funding her PhD studentship (CS). The work was also supported by funding from The Royal Society (Dorothy Hodgkin Fellowship DH160106) (RT).

